# Novel metric for hyperbolic phylogenetic tree embeddings

**DOI:** 10.1101/2020.10.09.334243

**Authors:** Hirotaka Matsumoto, Takahiro Mimori, Tsukasa Fukunaga

## Abstract

Advances in experimental technologies such as DNA sequencing have opened up new avenues for the applications of phylogenetic methods to various fields beyond their traditional application in evolutionary investigations, extending to the fields of development, differentiation, cancer genomics, and immunogenomics. Thus, the importance of phylogenetic methods is increasingly being recognized, and the development of a novel phylogenetic approach can contribute to several areas of research. Recently, the use of hyperbolic geometry has attracted attention in artificial intelligence research. Hyperbolic space can better represent a hierarchical structure compared to Euclidean space, and can therefore be useful for describing and analyzing a phylogenetic tree. In this study, we developed a novel metric that considers the characteristics of a phylogenetic tree for representation in hyperbolic space. We compared the performance of the proposed hyperbolic embeddings, general hyperbolic embeddings, and Euclidean embeddings, and confirmed that our method could be used to more precisely reconstruct evolutionary distance. We also demonstrate that our approach is useful for predicting the nearest-neighbor node in a partial phylogenetic tree with missing nodes. This study highlights the utility of adopting a geometric approach for further advancing the applications of phylogenetic methods.

The demo code is attached as a supplementary file in a compiled jupyter notebook. The code used for analyses is available on GitHub at https://github.com/hmatsu1226/HyPhyTree.

## Introduction

The advancement of DNA sequencing technologies has enabled the determination of the genome sequences of different organisms, resulting in the reconstruction of various types of gene and species trees (1–3). Phylogenetic analyses based on these trees have contributed substantially to research fields such as the identification of gene–gene associations and gene functions (4), relationships between evolution and disease (5), evolutionary dynamics of pathogens (6–9), and bacterial taxonomy (10). However, there is increasing recognition of the necessity for developing novel computational phylogenetic methods to further accelerate the application of phylogenetic analyses in various researches (11–13).

Furthermore, recent advances in lineage-tracing methods based on single-cell and genome-editing technologies have led to elucidation of cellular lineages (14), and phylogenetic methods have played an essential role in reconstructing and analyzing these cellular lineages. In addition, phylogenetic methods have contributed to research in cancer genomics (evolution of cancer) (15) and immunogenomics (evolution of antibody lineages) (16, 17). In addition to the essential role of phylogenetic methods in evolutionary biology, these approaches are increasingly becoming relevant for other research areas; thus, the development of novel phylogenetic methods can contribute broadly to various fields in the life and biomedical sciences.

Hyperbolic geometry, as a non-Euclidean form of geometry, has attracted attention in artificial intelligence research, with several methods using hyperbolic geometry recently developed for various applications. Rerepresentation learning models have shown that hyperbolic space could be used to represent latent hierarchical structures, exhibiting significant performance improvements over other approaches (18). This finding motivated several subsequent machine-learning studies using hyperbolic space, including the construction of hyperbolic neural networks (19). In addition, a novel approach for hierarchical clustering that optimizes the coordinates in hyperbolic space was proposed (20).

The coordinates in two-dimensional hyperbolic geometry can be described with the Poincaré disk model (or the Poincaré ball model for three-dimensional or *n*-dimensional hyperbolic space). In contrast to the Euclidean space, where the shortest path between two points (i.e., a geodesic) is a straight line, a geodesic in the Poincaré disk is an arc (Fig. 1A). This characteristic of geodesics on the Poincaré disk effectively represents a hierarchical structure because the geodesic provides a better match to a tree structure than the path on Euclidean space represented as a straight line (Fig. 1B). In addition, the area of the Poincaré disk grows exponentially in accordance with the distance from the origin, which is an advantageous characteristic for representing the nodes of a tree that increase in number exponentially with branching of the tree (Fig. 1C). These characteristics of a Poincaré disk are effective for embedding a phylogenetic tree, and visualization tools based on such hyperbolic phylogenetic tree embeddings have been developed (21, 22). Hyperbolic embeddings have also recently been used to reconstruct cell lineage trees from single-cell RNA-sequencing data (23, 24). In addition, the hyperbolic space has been used for several other types of biological studies such as for analyzing protein function based on protein interaction network embedding (25) and interpreting the mechanisms of olfactory space (26). Since hyperbolic embeddings are effective for analyzing biological data with a hierarchical structure, they are expected to be useful for various types of phylogenetic methods. To extend the applications of hyperbolic space for phylogenetic methods, we here propose a novel metric for accurate phylogenetic tree embedding in hyperbolic space. The proposed method is expected to contribute to the development of novel phylogenetic methods using hyperbolic space.

**Fig. 1.**
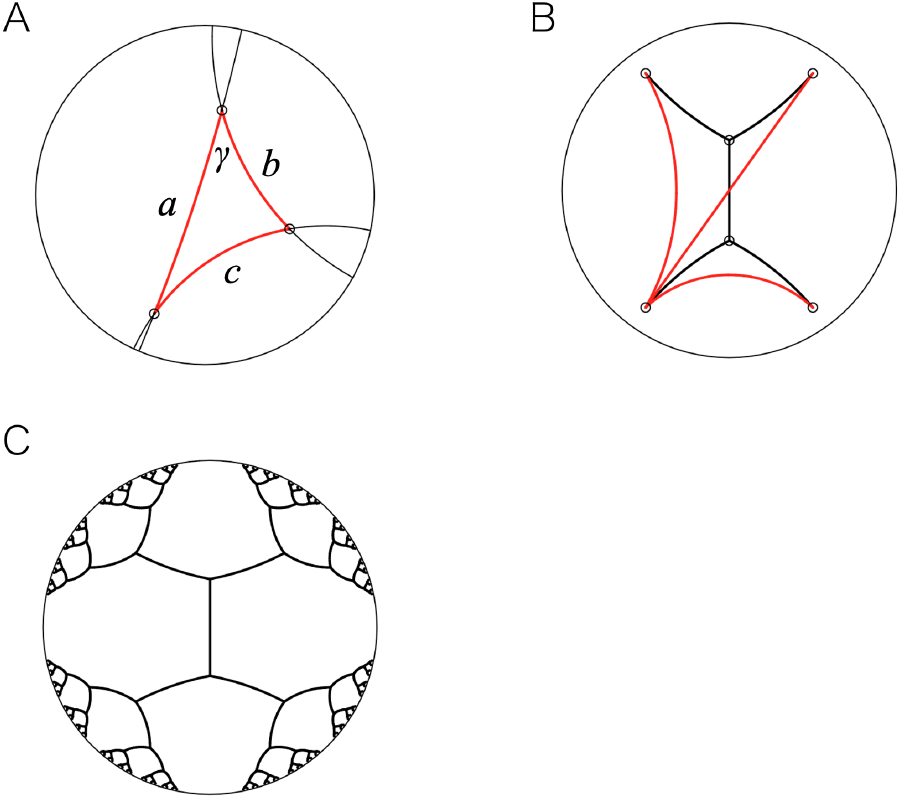
Graphical schematic of the Poincaré disk. (A) Geodesics and triangle on the Poincaré disk. (B) Geodesics for the external nodes of a simple tree structure. (C) Tree embedding on the Poincaré disk. The geodesic distances based on the Poincaré disk for all branches are equal.

For both Euclidean and hyperbolic embeddings, the input data are generally represented as a distance matrix among data points, and objective functions are designed to preserve the input distances with the geodesic distances on the embedded space. In the case of phylogenetic tree embeddings, the distance between two nodes on the tree corresponds to the evolutionary distance, which is the sum of branch lengths between two nodes (additivity). For example, considering the two external nodes *i* and *j* and their most recent common ancestor (MRCA) node *k*, the evolutionary distance *d_ij_* represents the sum of the respective distance of each node to the MRCA, *d_ik_* + *d_jk_*. Although hyperbolic phylogenetic tree embedding is expected to preserve distance information better than Euclidean phylogenetic tree embedding, a general embedding approach still cannot perfectly preserve the evolutionary distances of the phylogenetic tree, even when using hyperbolic space. This is because the additivity can only accurately reflect the geodesic distance when all nodes are located on a single geodesic line, and this is not the case for tree structures constructed with general embeddings. To overcome this limitation, we developed a novel metric that allows for embedding the evolutionary distance in a more precise manner. Specifically, the proposed metric is based on the cosine rule in hyperbolic geometry (i.e., the hyperbolic law of cosines), and the evolutionary distance is transformed based on our metric prior to incorporating the hyperbolic embeddings. We compared the performance of conventional general Euclidean and hyperbolic embeddings with that of our proposed embeddings, which confirmed that our approach could precisely reconstruct the evolutionary distance in a variety of scenarios. The proposed embeddings also exhibit an advantageous property in that the angles among external nodes and their MRCA node are well controlled, which would be useful for analyses including angle information. In addition, we investigated the ability of the three types of embeddings to predict the nearest node of an external node not included in the partial phylogenetic tree, demonstrating that our proposed approach had the best predictive performance. These results demonstrate the effectiveness of our metric for phylogenetic embeddings and the potential of adopting a geometric approach for phylogenetic analyses. The importance of novel phylogenetic methods has been increasingly recognized to handle the new and abundant data emerging from the current genomic era. In this regard, our approach has potential to contribute to several types of phylogenetic analyses by proposing a novel concept of “tree thinking” (27) based on geometric thinking.

## Material and methods

### Novel metric for hyperbolic phylogenetic tree embeddings

The novel metric developed was based on the hyperbolic law of cosines to represent the evolutionary distance precisely in hyperbolic space. For a hyperbolic triangle with side lengths *a, b*, and *c*, and angle (in radians) *γ* (see Fig. 1A), the hyperbolic law of cosines is satisfied, as follows:

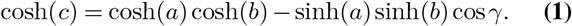

In the case of *γ* = *π*/2, the above equation becomes

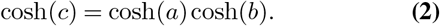

For the two nodes *i* and *j* in a phylogenetic tree and MRCA node *k*, we describe the evolutionary distances between any two nodes (*i, j*), (*i, k*), and (*j, k*) as *d_ij_, d_ik_*, and *d_jk_*, respectively. We also describe the geodesic distances in the hyperbolic space as 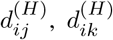, and 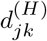. If we assume that the angle ∠*ikj* is *π*/2 and the geodesic distance satisfies 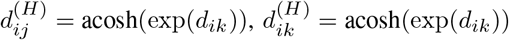, and 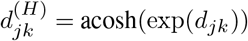, we can derive the following equation from Eq. 2:

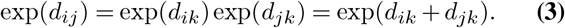

The above equation shows that the additivity of evolutionary distances (*d_ij_* = *d_ik_* + *d_jk_*) can be represented if the above assumptions are satisfied.

Thus, we translate the evolutionary distance between two nodes, *i* and *j*, as *X_ij_* = acosh(exp(*d_ij_*)) and embed the translated distance matrix *X* onto the hyperbolic space (Poincaré ball) instead of directly embedding the evolutionary distance matrix.

### Phylogenetic tree embeddings

The evolutionary distance matrix is represented as *D* (with *D_ij_* considered to be the evolutionary distance *d_ij_*), and then *D* was embedded with both Euclidean and general hyperbolic embeddings, in addition to hyperbolic embeddings with the proposed metric (Fig. 2). The performance of each approach was then compared.

**Fig. 2.**
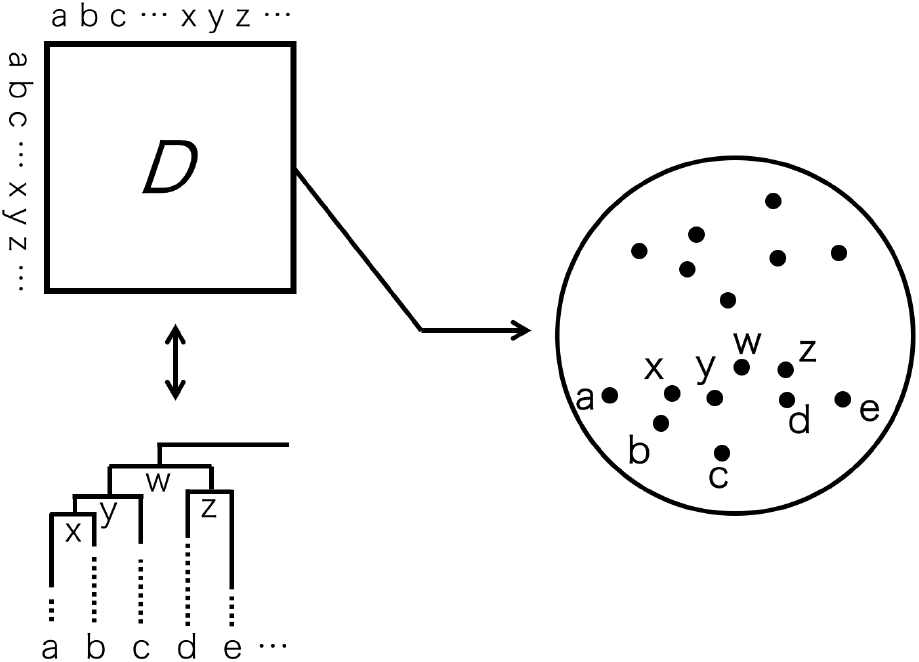
Graphical representation of hyperbolic phylogenetic tree embeddings. The input evolutionary distance matrix *D* contains all nodes, including the internal nodes of the phylogenetic tree. In our proposed embeddings, the input matrix is the translated matrix *X*, instead of *D*.

For Euclidean embeddings, we used Sammon mapping, which is a type of multidimensional scaling (MDS) (28). Toward this end, we used the *sammon* function in the MASS package in R language, and embedded *D* for the Euclidean space with various dimensions (*M*). The performance of the embeddings was evaluated by calculating the mean squared error (MSE) between *D_ij_* and the Euclidean distance *d_E_*(*z_i_, z_j_*), where *z_i_* represents the coordinate of node *i* of the embedded space.

For general hyperbolic embeddings, we used the *hydraPlus* function in the hydra package in R (29), and embedded *D* with parameters *curvature* = 1 and *alpha* = 1 under various dimensions *M*. The performance was then evaluated by calculating the MSE between *D_ij_* and the geodesic distance on the Poincaré ball *d_P_*(*z_i_, z_j_*), where the distance is defined as follows:

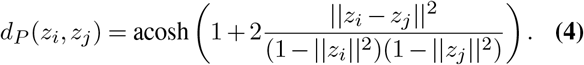

We also used the *hydraPlus* function for hyperbolic embeddings with the proposed metric. We embedded the translated matrix *X*, where *X_ij_* = acosh(exp(*D_ij_*)), and evaluated its performance by calculating the MSE between *D_ij_* and the inverse transformed distance log(cosh(*d_P_*(*z_i_, z_j_*))).

### Folding-in internal node

In the previous section, we considered the distance matrix *D* for all nodes in a phylogenetic tree as the input, including internal nodes, and embedded all nodes simultaneously. We further evaluated the performance of each type of embedding in the case of “folding-in” the internal nodes. In this case, the input was a distance matrix for external nodes, and the external nodes were first embedded, followed by optimization of the coordinates of the internal nodes for the embedding space (Fig. 3).

**Fig. 3.**
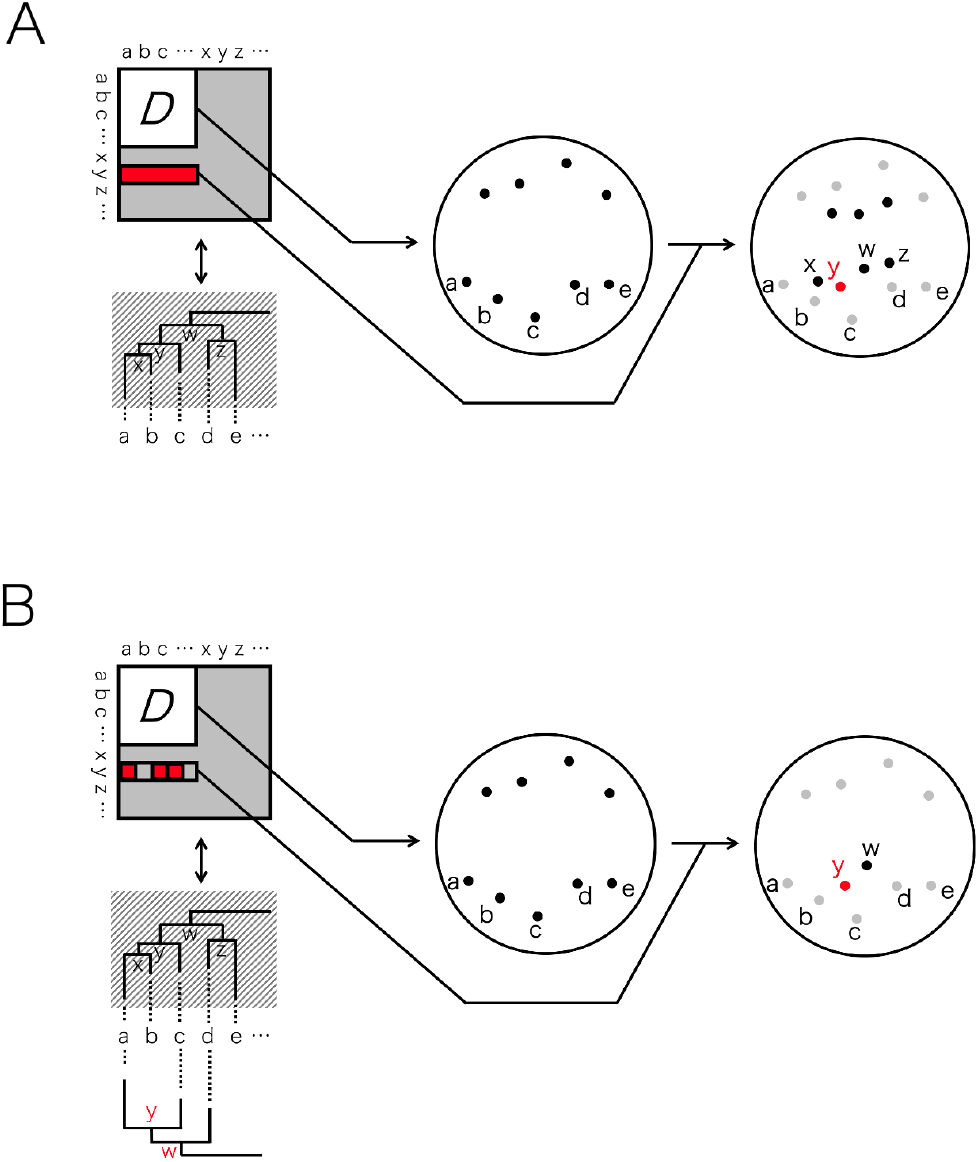
Graphical representation of the hyperbolic embeddings of a distance matrix corresponding to the external nodes and folding-in of the internal nodes of the complete phylogenetic tree (A) and partial tree (B).

We first conducted these steps for complete phylogenetic tree. The external nodes were first embedded, and the coordinates of each internal node is optimized numerically and independently (Fig. 3A). The objective function used for optimizing the coordinates of an internal node was the minimization of the MSE between the true evolutionary distances and the reconstructed evolutionary distance based on the geodesic distance for the internal node and all external nodes.

We next embedded the internal nodes of a partial tree, which is the phylogenetic tree for only a portion of the external nodes (Fig. 3B). The objective function used here is the minimization of the MSE between the true evolutionary distances and the reconstructed evolutionary distance for the internal node and external nodes included in the partial tree.

### Prediction of the MRCA

Based on the results of each method of embedding, we developed a new indicator to predict the MRCA of two external nodes *i* and *j*. The indicator is based on the angle information on the embedded space, which was used to determine whether the internal node *k* is the MRCA of *i* and *j*. When node *k* is the MRCA for *i* and *j*, the additivity of the evolutionary distance (*d_ij_* = *d_ik_* + *d_jk_*) is satisfied. The additivity is also satisfied for the outgroup node *o*(*d_io_* = *d_ik_* + *d_ok_* and *d_jo_* = *d_jk_* + *d_ok_*). If the internal node *k*′ is a more distant (i.e., not the most recent) ancestor, then *d_ik′_* + *d_jk′_* becomes larger than *d_ij_*, and the angle on the embedded space will be ∠*ik′ j* < ∠*ikj*, where *k* is the MRCA of *i* and *j*. We calculated the angles ∠*ikj*, ∠*iko*, and ∠*jko* based on the embedded coordinates, and then used the minimum value of each angle as the “angle-score,” which served as the basis for predicting whether or not *k* is the MRCA.

We evaluated the performance of angle-score to predict the MRCA for each embedding type by randomly selecting the external nodes *i* and *j* 1000 times, and used *i, j*, and their MRCA *k* as the positive-control dataset. We also randomly selected the external node *l*, and used the combination *i, l*, and *k* as the negative-control dataset (if the MRCA of *i* and *l* was *k*, we resampled *l* randomly). We performed the above process for each random simulated tree and calculated the mean of the area under the receiver operating characteristic (ROC) curve (AUC) values to evaluate the prediction of the MRCA with the angle-score for each embedding method.

### Prediction of the nearest-neighbor node in the partial tree

Numerous genomes have been determined in the genome era, and the importance of integrating multiple partial phylogenetic trees (30, 31) or placing a novel species into an already-established phylogenetic tree (phylogenetic placement) (32) is increasing. Therefore, we evaluated the ability of our method to predict the nearest-neighbor node in the partial tree for some external nodes missing from the tree. First, we embedded the external nodes and folding-in the partial tree as described above (Fig. 3(B)). Second, we predicted the nearest-neighbor node in the partial tree for an external node that was not included the tree based on the geodesic distance for each embedding method. We evaluated the performance of each embedding by calculating the rank of the actual nearest-neighbor node calculated from the complete phylogenetic tree.

### Datasets

We randomly generated phylogenetic tree shapes comprising 100 external nodes using the *rtree* function (33). We determined each branch length with *α*(1 – log(*u*(*e* – 1) + 1)), where *u* is a uniform random value and *α* corresponds to the scaling factor, which was set to values of 0.25, 0.5, and 1 for this analysis. We added another external node as an outgroup for the phylogenetic tree, resulting in a tree with 101 external nodes. We generated the trees 100 times independently for each *α* value. For analyses of the partial trees, we randomly selected 20 external nodes in addition to the outgroup node and extracted the partial tree corresponding to these external nodes set from the complete phylogenetic trees. Hereafter, we refer to the dataset with *α* = 0.25, 0.5, and 1.0 as Data_.25_, Data_.5_, and Data_1_, respectively.

We also used the phylogenetic tree of primates downloaded from TimeTree(34) for evaluation. We used the genus-level phylogenetic tree and added one outgroup node, resulting in a tree with 77 external nodes. First, we normalized the branch lengths such that the maximum branch length was 1.0, and then perturbed the branch lengths by adding 0.5(1 – log(*u*(*e* – 1) +1)). We generated the branch length-perturbed trees 100 times independently. The partial trees were generated in the same manner as implemented for the random phylogenetic trees. We refer to the dataset as Data_pri_.

## Results

### Full node embeddings

We embedded the evolutionary distance matrix *D*, including the internal nodes, with each embedding method, and evaluated the performance of each method according to the MSE calculated between the actual evolutionary distance and reconstructed evolutionary distance based on the coordinate of the embedded space. The mean MSE values for the dataset Data_.5_ of MDS, general hyperbolic embeddings (H1), and the proposed hyperbolic embeddings (H2) for various embedding dimensions *M* are shown in Table 1. The results for the dataset Data_.25_, Data_1_, and Data_pri_ are provided in the supplementary note.

**Table 1.** Mean MSE values of full node embeddings forthe dataset Data_.5_ for each embedding method (columns): Euclidean embeddings (MDS), general hyperbolic embeddings (H1), and the proposed hyperbolic embeddings (H2). The rows represent the dimensions of the embeddings.

| | $M=5$ | $M=10$ | $M=20$ | $M=30$ |
| --- | --- | --- | --- | --- |
| MDS | $1.4 \times 10^{-1}$ | $4.4 \times 10^{-2}$ | $2.3 \times 10^{-2}$ | $2.0 \times 10^{-2}$ |
| H1 | $1.4 \times 10^{-2}$ | $4.3 \times 10^{-3}$ | $3.2 \times 10^{-3}$ | $3.2 \times 10^{-3}$ |
| H2 | $2.3 \times 10^{-2}$ | $4.0 \times 10^{-3}$ | $5.6 \times 10^{-4}$ | $1.6 \times 10^{-4}$ |

The mean MSE values of MDS were larger than those of H1 and H2 for all cases, and the hyperbolic space offered a better representation of the phylogenetic trees. Although the mean MSE value of H1 was the smallest for *M* = 5, the values of H2 were the smallest for *M* ≥ 10. In particular, the mean MSE value of H2 was less than one-tenth the value of H1 for *M* = 30.

We also investigated the angles (degrees) of any external nodes *i* and *j* and their MRCA *k*(∠*ikj*) for each embedding method under various *M* values (Fig. 4). The dispersion of angles of MDS and H1 were large, even when the dimension *M* was increased. In contrast, all of the angles of H2 merged to approximately 90° in accordance with increasing *M*. The median angle of H2 for *M* = 5 was over 100°, which had a gap to satisfy Eq. 2, explaining the larger MSE value of H2 than that of H1 for *M* = 5.

**Fig. 4.**
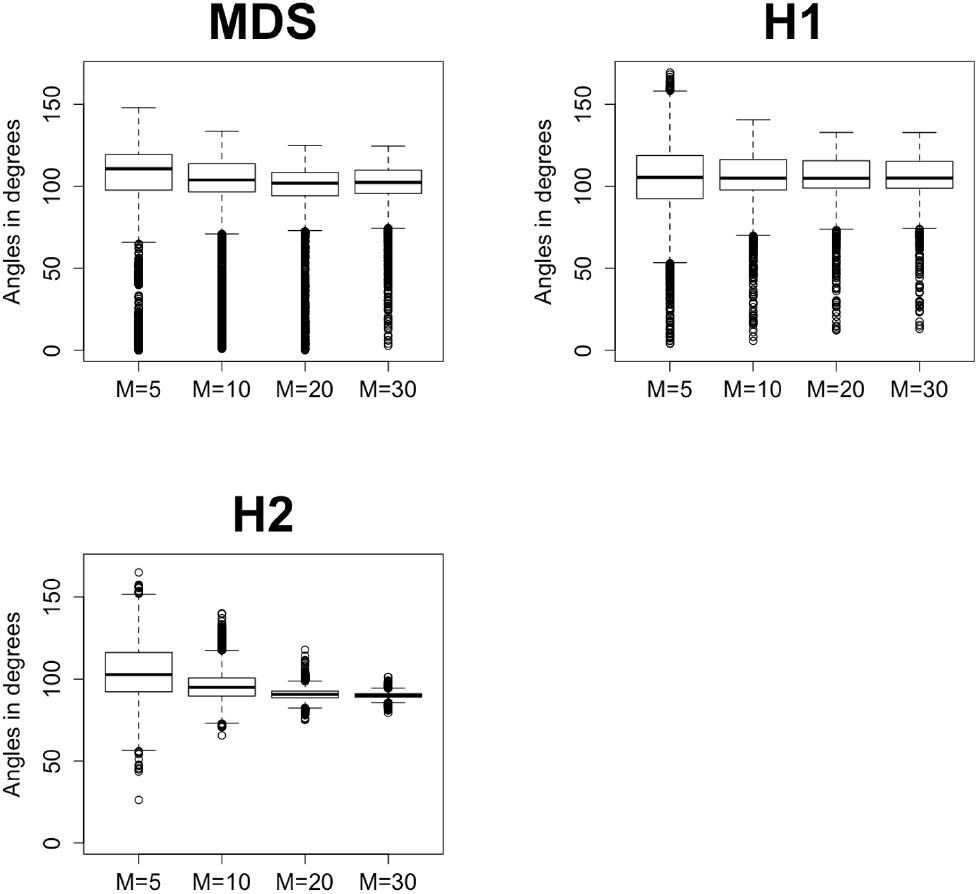
Angles (degrees) of two external nodes and their MRCA for the dataset Data_.5_ for each embedding method and each dimension.

Thus, the proposed hyperbolic embeddings can precisely represent the phylogenetic tree, especially with large embedding dimensions. In addition, our proposed embeddings offer an advantage with respect to the angle, which will be useful for analyses or machine learning using angle information.

### Prediction of the MRCA

Based on the embeddings described in the subsection above, we next evaluated the ability of each method to predict the MRCA based on the angle information. We calculated the angle-score for the positive and negative datasets (see Methods section), and evaluated the performance according to the AUC values of ROC curves for each embedding method. The mean AUC values of 100 simulated phylogenetic trees with *α* = 0.5 (Data_.5_) are shown in Table 2.

**Table 2.** Mean AUC values for the ability to identify the MRCA based on the anglescore for the dataset Data_.5_ for each embedding method (columns). The rows represent the dimensions of the embeddings.

| | $M=5$ | $M=10$ | $M=20$ | $M=30$ |
| --- | --- | --- | --- | --- |
| MDS | 0.62 | 0.71 | 0.78 | 0.79 |
| H1 | 0.80 | 0.87 | 0.90 | 0.90 |
| H2 | **0.91** | **0.95** | **0.97** | **0.99** |

The AUC values of the proposed method (H2) were the highest compared with those of other embedding methods for all dimensions *M*. In particular, our method could predict the MRCA almost perfectly for *M* = 30. Interestingly, the AUC value of H2 was significantly higher than that of H1, despite the fact that the MSE for standard hyperbolic embeddings was better than that of our embedding method with *M* = 5. This result implies that the proposed embedding method maintains the integrity of the angles even when the dimensions of the embeddings are small. This tendency was consistent for other datasets (see supplementary note).

### External node embeddings and folding-in internal nodes

In the previous analysis, we embedded all nodes, including the internal nodes, of a phylogenetic tree simultaneously. Next, we embedded the external nodes and then appended the internal nodes onto the embedding space (Fig. 3A). The mean MSE values for the dataset Data_.5_ for each embedding method are shown in Table 3.

**Table 3.** Mean MSE values of external node embeddings and folding-in internal nodes for each embedding method (columns). Rows represent the dimensions of the embeddings.

| | $M=5$ | $M=10$ | $M=20$ | $M=30$ |
| --- | --- | --- | --- | --- |
| MDS | $1.0 \times 10^{-1}$ | $4.2 \times 10^{-2}$ | $2.2 \times 10^{-2}$ | $2.0 \times 10^{-2}$ |
| H1 | **$1.4 \times 10^{-2}$** | $4.6 \times 10^{-3}$ | $3.6 \times 10^{-3}$ | $3.5 \times 10^{-3}$ |
| H2 | $2.3 \times 10^{-2}$ | **$4.0 \times 10^{-3}$** | **$6.2 \times 10^{-4}$** | **$2.3 \times 10^{-4}$** |

Similar to the previous results, the mean MSE values of the proposed embedding method were better than those of other embeddings with folding-in of internal nodes, especially for *M* = 30. These results indicate that the proposed embeddings can be trained to learn the appropriate space even if the only information available is the evolutionary distance of external nodes.

### External node embeddings and folding-in internal nodes of the partial tree

We further evaluated the performance of each embedding method when the evolutionary distances of some nodes are only partially known. As an example, we first embedded the external nodes and then appended the internal nodes of the partial tree according to the evolutionary distances for the external nodes included in the tree (Figure 3B). Finally, we calculated the MSE for the evolutionary distance between the external nodes that were not included in the tree and the internal nodes. The mean MSE values for the dataset Data_.5_ for each embedding method are shown in Table 4. Similar to the previous results, the mean MSE values of the proposed embeddings were better than those of other embeddings, especially for *M* = 30. Thus, the proposed method can also effectively embed nodes when there is missing data on the evolutionary distances for some nodes. This approach can be useful for analyses of phylogenetic trees with only partial external nodes available.

**Table 4.**
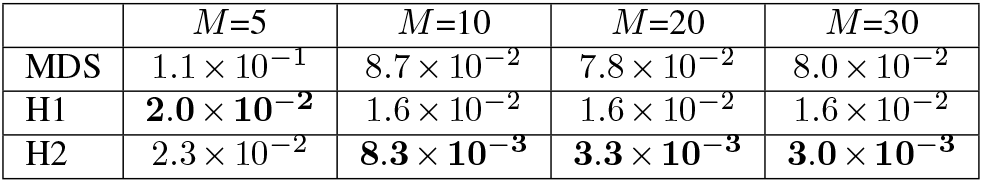
Mean MSE values of external node embeddings and folding-in internal nodes of the partial tree for each embedding method (columns). The rows represent the dimensions of the embeddings.

### Prediction of nearest-neighbor nodes in the partial tree

Based on the embeddings for the partial tree, we investigated the ability to predict the nearest-neighbor node for an external node that is not included in the partial tree. The prediction performances of the different embedding methods were compared by calculating the rank of the actual nearest-neighbor node for the external nodes based on the respective geodesic distances. The actual nearest-neighbor node can be determined from the complete phylogenetic tree. The mean ranks of each method for the dataset Data_.5_ are shown in Table 5, and the results for other datasets are shown in the supplementary note.

**Table 5.** Mean rank of predicting the true nearest-neighbor nodes in the partial tree for external nodes not included the tree. The rank is based on the geodesic distances for each embedding method.

| | $M=5$ | $M=10$ | $M=20$ | $M=30$ |
| --- | --- | --- | --- | --- |
| MDS | 3.07 | 2.47 | 2.13 | 2.04 |
| H1 | 1.75 | 1.52 | 1.47 | 1.46 |
| H2 | **1.43** | **1.24** | **1.18** | **1.14** |

**Table 6.** Mean MSE values of full node embeddings for the dataset Data_.25_, Data_1_, and Data_pri_.

| | $M=5$ | $M=10$ | $M=20$ | $M=30$ |
| --- | --- | --- | --- | --- |
| MDS | $3.9 \times 10^{-2}$ | $1.3 \times 10^{-2}$ | $6.1 \times 10^{-3}$ | $5.5 \times 10^{-3}$ |
| H1 | **$7.7 \times 10^{-3}$** | **$2.7 \times 10^{-3}$** | $2.0 \times 10^{-3}$ | $2.0 \times 10^{-3}$ |
| H2 | $2.0 \times 10^{-2}$ | $4.5 \times 10^{-3}$ | **$8.4 \times 10^{-4}$** | **$2.8 \times 10^{-4}$** |
| MDS | $5.1 \times 10^{-1}$ | $1.6 \times 10^{-1}$ | $8.9 \times 10^{-2}$ | $7.8 \times 10^{-2}$ |
| H1 | **$1.0 \times 10^{-2}$** | $3.7 \times 10^{-3}$ | $3.3 \times 10^{-3}$ | $3.2 \times 10^{-3}$ |
| H2 | $1.2 \times 10^{-2}$ | **$1.3 \times 10^{-3}$** | **$1.2 \times 10^{-4}$** | **$2.9 \times 10^{-5}$** |
| MDS | $3.0 \times 10^{-1}$ | $1.0 \times 10^{-1}$ | $4.4 \times 10^{-2}$ | $3.8 \times 10^{-2}$ |
| H1 | **$9.3 \times 10^{-3}$** | $3.7 \times 10^{-3}$ | $3.3 \times 10^{-3}$ | $3.3 \times 10^{-3}$ |
| H2 | $1.2 \times 10^{-2}$ | **$1.5 \times 10^{-3}$** | **$1.6 \times 10^{-4}$** | **$3.7 \times 10^{-5}$** |

**Table 7.** Mean AUC values in identifying the MRCA for the dataset Data_.25_, Data_1_, and Data_pri_.

| | $M=5$ | $M=10$ | $M=20$ | $M=30$ |
| --- | --- | --- | --- | --- |
| MDS | 0.64 | 0.73 | 0.76 | 0.76 |
| H1 | 0.68 | 0.75 | 0.80 | 0.80 |
| H2 | **0.81** | **0.91** | **0.94** | **0.96** |
| MDS | 0.65 | 0.79 | 0.76 | 0.82 |
| H1 | 0.93 | 0.95 | 0.96 | 0.96 |
| H2 | **0.96** | **0.98** | **0.99** | **1.00** |
| MDS | 0.68 | 0.79 | 0.89 | 0.90 |
| H1 | 0.98 | 0.99 | **1.00** | **1.00** |
| H2 | **1.00** | **1.00** | **1.00** | **1.00** |

**Table 8.** Mean rank of true nearest-neighbor nodes in the partial trees for the external nodes not included the trees for the dataset Data_.25_, Data_1_, and Data_pri_.

| | $M=5$ | $M=10$ | $M=20$ | $M=30$ |
| --- | --- | --- | --- | --- |
| MDS | 3.20 | 2.40 | 2.08 | 1.99 |
| H1 | 2.17 | 1.82 | 1.71 | 1.70 |
| H2 | **1.71** | **1.37** | **1.27** | **1.19** |
| MDS | 3.02 | 2.42 | 2.05 | 1.97 |
| H1 | 1.29 | 1.23 | 1.22 | 1.21 |
| H2 | **1.22** | **1.16** | **1.12** | **1.11** |
| MDS | 3.40 | 2.65 | 2.35 | 2.24 |
| H1 | 1.42 | 1.28 | 1.26 | 1.26 |
| H2 | **1.24** | **1.15** | **1.13** | **1.13** |

The mean ranks using embeddings based on hyperbolic space (H1 and H2) were superior to those obtained using Euclidean embeddings. Moreover, the mean ranks of the proposed method demonstrated the best performance overall under all conditions. In particular, the mean rank of our method was close to 1 with *M* = 30. Therefore, our approach would be effective for analyzing a partial tree and assigning missing nodes.

## Discussion

Although numerous algorithms for phylogenetic tree reconstruction and phylogenetic analyses have been developed with superior performance, there has been limited discussion of these algorithms from the perspective of geometry. Our results demonstrate that a geometric view has potential to provide novel knowledge for rethinking and improving these algorithms. As an example, we here focus on the application of such a geometric perspective for improving the integration of multiple gene trees (35).

The coordinates of the *M*-dimensional Poincaré ball model *z_i_* can be transformed to the coordinates of the *M*-dimensional hyperbolic geometry *x_i_* = {*x_i,k_*|*k* = 1,…, *M* + 1}, where *x_i_* satisfies 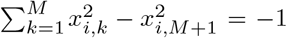. The geodesic distance based on the above coordinates (hyperbolic geometry version of Eq. 4) is then defined as follows:

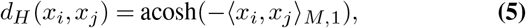

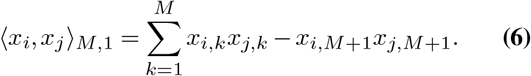

Using our novel metric for the conversion of evolutionary distance, acosh(exp(*d_ij_*)), the pseudo-inner product (〈*x_i_, x_j_*〉_*M*,1_) has the following relationship to evolutionary distance:

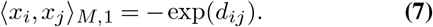

Here, we consider two genes, *a* and *b*, and their phylogenetic gene trees independently. The evolutionary distances for each tree are then 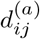 and 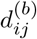, respectively. Based on Eq. 7, the mean of the pseudo-inner product satisfies the following relationship:

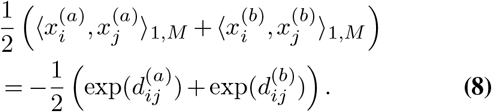

Then, we define a new vector, 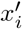, which is combined with 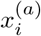 and 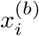 as follows:

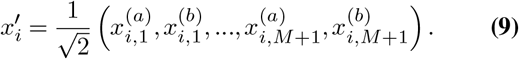

Using the above vector, the mean of the pseudo-inner product can be represented with the following bilinear form:

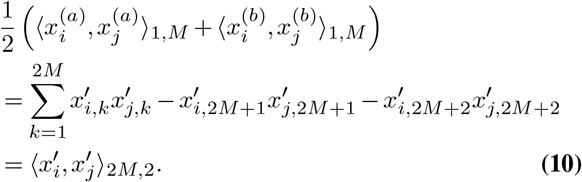

This form is known as an inner product related to a pseudo-Euclidean space. Therefore, the pseudo-Euclidean space may be useful for representing the ensembled trees. Furthermore, our approach suggests that the arithmetic mean of the exponential of the evolutionary distance is useful for creating an ensemble of multiple gene trees.

Recently, novel kernels that can deal with indefinite inner products have been developed in representation learning studies (36, 37). The inner product of Eq. 10 is associated with these kernels, and we plan to extend our phylogenetic approach by adopting these techniques in future work.

In conclusion, we here propose and validate a novel metric for hyperbolic phylogenetic tree embeddings, which could precisely reconstruct evolutionary distance. Our method had a particular advantage over other embedding methods in that the angles were consistent for any two external nodes and the node of their MRCA. Moreover, our approach was shown to be useful for predicting the MRCA and the nearest-neighbor node in a partial tree with missing nodes. These results highlight the possibility of applying “geometric-thinking” as an effective approach to the novel “tree-thinking” approach. This is particularly relevant as evolutionary analyses are expanding to different research fields beyond evolutionary inferences themselves, including differentiation, immunogenomics, and cancer evolution, requiring a novel phylogenetic strategy, and we will extend our approach to these analyses in future work.

## ACKNOWLEDGEMENTS

This work was supported by the Japan Society for the Promotion of Science [grant numbers JP20H05582]. We would like to thank Haru Oono Negami and Dr. Yasuhiro Kojima for providing helpful advice about the use of hyperbolic geometry. We would also like to thank Mr. Akihiro Matsushima and Mr. Manabu Ishii at the Laboratory for Bioinformatics Research, RIKEN BDR, for their assistance with the IT infrastructure for the data analysis.

## Supplementary Note 1: Evaluation with various datasets

We evaluated full node embeddings for the datasets Data_.25_, Data_1_, and Data_pri_, respectively. The mean MSE values for each embedding method are shown in Table 6. Similar to the results described in the main text for the dataset Data_.5_, the mean MSE values of the proposed embedding method were significantly better than those of other embedding methods for both *M* = 20 and 30. We also evaluated the MSE values of folding-in internal nodes and the folding-in partial tree, which showed the same tendency of superiority of the proposed method for all datasets, consistent with the results described in the main text.

In addition, the proposed method showed the highest performance in predicting the MRCA based on the angle-score (Table 7) and in predicting the nearest-neighbor node in the partial tree (Table 8) for all datasets.

