## supplementary file for "Novel metric for hyperbolic phylogenetic tree embeddings"

demo


In [22]:

```
library(hydra)
library(ape)
library(MASS)
library(stringr)
```

In [23]:

```
#generate simulation tree

Nleaves <- 100
alpha <- 0.5

tree <- rtree(Nleaves)
tree$tip.label <- 1:Nleaves
for(j in 1:length(tree$edge.length)){
    tree$edge.length[j] <- alpha * (1 - log(runif(1)*(exp(1)-1)+1))
}

tmp <- paste("(",str_sub(write.tree(tree), end=-2),":0.1,",Nleaves+1,":2);",sep="")
tree <- read.tree(text=tmp)

plot(tree, type="unrooted")
```

In [28]:

```
#geodesic distance on Poincare ball
Poincare_dist <- function(u,v){
    if(sum(u**2)>=1 || sum(v**2)>=1){
        return(Inf)
    }else{
        return(acosh(1+2*(sum((u-v)**2))/(1-sum(u**2))/(1-sum(v**2))))
    }
}

#embedding dimension
Ms <- c(5,10,20,30)

Nleaves <- length(tree$tip.label)
tree$mrca <- mrca(tree)
D <- dist.nodes(tree)

Nall <- dim(D)[1]
X1 <- D
X2 <- acosh(exp(D))

MSE_H1 <- rep(0, length(Ms))
MSE_H2 <- rep(0, length(Ms))
MSE_MDS <- rep(0, length(Ms))

angle_H1 <- matrix(0, Nleaves*(Nleaves-1)/2, length(Ms))
angle_H2 <- matrix(0, Nleaves*(Nleaves-1)/2, length(Ms))
angle_MDS <- matrix(0, Nleaves*(Nleaves-1)/2, length(Ms))

for(M in Ms){
    #general hyperbolic embeddings
    X1.hydra <- hydraPlus(X1, dim=M, curvature=1, alpha=1, equi.adj=0, control=list(return.dist=1, isotropic.adj=FALSE))
    Z1 <- X1.hydra$r * X1.hydra$directional
    X1.hydra$dist <- hydra:::get.distance(X1.hydra$r, X1.hydra$directional, X1.hydra$curvature)
    
    #our hyperbolic embeddings
    X2.hydra <- hydraPlus(X2, dim=M, curvature=1, alpha=1, equi.adj=0, control=list(return.dist=1, isotropic.adj=FALSE))
    Z2 <- X2.hydra$r * X2.hydra$directional
    X2.hydra$dist <- hydra:::get.distance(X2.hydra$r, X2.hydra$directional, X2.hydra$curvature)

    #Euclidean embeddings
    X1.mds <- sammon(X1, k=M)
    X1.mds$dist <- matrix(0, Nall, Nall)
    for(i in 1:Nall){
        for(j in 1:Nall){
            X1.mds$dist[i,j] <- sqrt(sum((X1.mds$points[i,]-X1.mds$points[j,])**2))
        }
    }

    #MSE
    MSE_H1[which(Ms==M)] <- sum((c(D[upper.tri(D)])-c(X1.hydra$dist[upper.tri(D)]))**2) / choose(Nall,2)
    MSE_H2[which(Ms==M)] <- sum((c(D[upper.tri(D)])-c(log(cosh(X2.hydra$dist[upper.tri(D)]))))**2) / choose(Nall,2)
    MSE_MDS[which(Ms==M)] <- sum((c(D[upper.tri(D)])-c(X1.mds$dist[upper.tri(D)]))**2) / choose(Nall,2)
    
    #Angle information
    idx <- 1
    for(i in 1:(Nleaves-1)){
        for(j in (i+1):Nleaves){
            k <- tree$mrca[i,j]
            
            #general hyperbolic embeddings
            a <- Poincare_dist(Z1[i,], Z1[k,])
            b <- Poincare_dist(Z1[j,], Z1[k,])
            c <- Poincare_dist(Z1[i,], Z1[j,])
            angle_H1[idx,which(Ms==M)] <- acos((cosh(a)*cosh(b)-cosh(c))/(sinh(a)*sinh(b)))
            
            #our hyperbolic embeddings
            a <- Poincare_dist(Z2[i,], Z2[k,])
            b <- Poincare_dist(Z2[j,], Z2[k,])
            c <- Poincare_dist(Z2[i,], Z2[j,])
            angle_H2[idx,which(Ms==M)] <- acos((cosh(a)*cosh(b)-cosh(c))/(sinh(a)*sinh(b)))
            
            #Euclidean embeddings
            a <- sqrt(sum((X1.mds$points[i,]-X1.mds$points[k,])**2))
            b <- sqrt(sum((X1.mds$points[j,]-X1.mds$points[k,])**2))
            c <- sqrt(sum((X1.mds$points[i,]-X1.mds$points[j,])**2))
            angle_MDS[idx,which(Ms==M)] <- acos((a**2+b**2-c**2)/(2*a*b))
            idx <- idx + 1
        }
    }
}
```

```
iter   10 value 469.924124
iter   20 value 359.854851
iter   30 value 337.300439
iter   40 value 326.905996
iter   50 value 321.835280
iter   60 value 319.830972
iter   70 value 318.984499
iter   80 value 318.574085
iter   90 value 318.380862
iter  100 value 318.223363
iter  110 value 318.079644
iter  120 value 317.845873
iter  130 value 317.715524
iter  140 value 317.677117
iter  150 value 317.659520
iter  160 value 317.644190
iter  170 value 317.622592
iter  180 value 317.598102
iter  190 value 317.579651
iter  200 value 317.561339
iter  210 value 317.548942
iter  220 value 317.542109
iter  230 value 317.537877
iter  240 value 317.535162
iter  250 value 317.533473
iter  260 value 317.532457
iter  270 value 317.531854
iter  280 value 317.531481
iter  290 value 317.531134
iter  300 value 317.530904
iter  310 value 317.530726
iter  320 value 317.530597
iter  330 value 317.530516
iter  340 value 317.530455
iter  350 value 317.530404
iter  360 value 317.530371
final  value 317.530365 
converged
iter   10 value 773.148977
iter   20 value 612.684388
iter   30 value 567.244804
iter   40 value 555.196951
iter   50 value 550.410432
iter   60 value 547.037610
iter   70 value 544.269346
iter   80 value 542.336828
iter   90 value 540.951704
iter  100 value 540.078809
iter  110 value 539.352970
iter  120 value 538.963440
iter  130 value 538.641373
iter  140 value 538.395977
iter  150 value 538.175296
iter  160 value 538.027486
iter  170 value 537.929465
iter  180 value 537.829203
iter  190 value 537.712995
iter  200 value 537.579099
iter  210 value 537.466639
iter  220 value 537.405087
iter  230 value 537.350947
iter  240 value 537.206537
iter  250 value 537.082128
iter  260 value 537.043590
iter  270 value 537.013772
iter  280 value 536.997887
iter  290 value 536.991560
iter  300 value 536.988414
iter  310 value 536.986693
iter  320 value 536.985403
iter  330 value 536.984764
iter  340 value 536.984051
iter  350 value 536.983313
iter  360 value 536.982805
iter  370 value 536.982557
iter  380 value 536.982429
iter  390 value 536.982327
iter  400 value 536.982274
iter  410 value 536.982240
iter  420 value 536.982218
final  value 536.982202 
converged
Initial stress        : 0.04706
stress after   8 iters: 0.04397
iter   10 value 193.531995
iter   20 value 107.411260
iter   30 value 91.476666
iter   40 value 86.342021
iter   50 value 84.457838
iter   60 value 83.538212
iter   70 value 82.868569
iter   80 value 82.408237
iter   90 value 82.043877
iter  100 value 81.768770
iter  110 value 81.637081
iter  120 value 81.551849
iter  130 value 81.481404
iter  140 value 81.425606
iter  150 value 81.381036
iter  160 value 81.347391
iter  170 value 81.324513
iter  180 value 81.303563
iter  190 value 81.281931
iter  200 value 81.267453
iter  210 value 81.252898
iter  220 value 81.239266
iter  230 value 81.225919
iter  240 value 81.213520
iter  250 value 81.203945
iter  260 value 81.195488
iter  270 value 81.185788
iter  280 value 81.178359
iter  290 value 81.170675
iter  300 value 81.161505
iter  310 value 81.150123
iter  320 value 81.142498
iter  330 value 81.136341
iter  340 value 81.130355
iter  350 value 81.124529
iter  360 value 81.117192
iter  370 value 81.108881
iter  380 value 81.097254
iter  390 value 81.083688
iter  400 value 81.069425
iter  410 value 81.054696
iter  420 value 81.042273
iter  430 value 81.029565
iter  440 value 81.014682
iter  450 value 80.994288
iter  460 value 80.974510
iter  470 value 80.955421
iter  480 value 80.940942
iter  490 value 80.924972
iter  500 value 80.901507
iter  510 value 80.872085
iter  520 value 80.838089
iter  530 value 80.813360
iter  540 value 80.793186
iter  550 value 80.778098
iter  560 value 80.767143
iter  570 value 80.761723
iter  580 value 80.757318
iter  590 value 80.753965
iter  600 value 80.751495
iter  610 value 80.749768
iter  620 value 80.748361
iter  630 value 80.747192
iter  640 value 80.746159
iter  650 value 80.744930
iter  660 value 80.743423
iter  670 value 80.742262
iter  680 value 80.741225
iter  690 value 80.740446
iter  700 value 80.739455
iter  710 value 80.738532
iter  720 value 80.737509
iter  730 value 80.736520
iter  740 value 80.735547
iter  750 value 80.734828
iter  760 value 80.734268
iter  770 value 80.733882
iter  780 value 80.733539
iter  790 value 80.733190
iter  800 value 80.732941
iter  810 value 80.732670
iter  820 value 80.732419
iter  830 value 80.732159
iter  840 value 80.731952
iter  850 value 80.731801
iter  860 value 80.731703
iter  870 value 80.731640
iter  880 value 80.731597
iter  890 value 80.731559
iter  900 value 80.731517
iter  910 value 80.731485
iter  920 value 80.731456
iter  930 value 80.731436
iter  940 value 80.731417
iter  950 value 80.731395
iter  960 value 80.731376
iter  970 value 80.731358
iter  980 value 80.731337
iter  990 value 80.731318
iter 1000 value 80.731298
final  value 80.731296 
stopped after 1001 iterations
iter   10 value 230.791066
iter   20 value 115.774993
iter   30 value 95.585635
iter   40 value 88.426281
iter   50 value 85.773240
iter   60 value 84.002434
iter   70 value 82.661546
iter   80 value 82.008343
iter   90 value 81.579221
iter  100 value 81.157698
iter  110 value 80.861195
iter  120 value 80.687873
iter  130 value 80.535068
iter  140 value 80.408435
iter  150 value 80.319926
iter  160 value 80.241662
iter  170 value 80.181684
iter  180 value 80.138858
iter  190 value 80.100240
iter  200 value 80.061881
iter  210 value 80.020797
iter  220 value 79.976013
iter  230 value 79.918569
iter  240 value 79.856192
iter  250 value 79.799444
iter  260 value 79.758096
iter  270 value 79.723619
iter  280 value 79.696441
iter  290 value 79.674375
iter  300 value 79.658251
iter  310 value 79.646953
iter  320 value 79.634746
iter  330 value 79.625972
iter  340 value 79.617327
iter  350 value 79.610085
iter  360 value 79.602463
iter  370 value 79.593532
iter  380 value 79.581339
iter  390 value 79.569794
iter  400 value 79.558228
iter  410 value 79.546273
iter  420 value 79.532337
iter  430 value 79.522904
iter  440 value 79.515579
iter  450 value 79.506263
iter  460 value 79.496516
iter  470 value 79.488122
iter  480 value 79.479631
iter  490 value 79.472923
iter  500 value 79.465471
iter  510 value 79.458366
iter  520 value 79.451069
iter  530 value 79.445244
iter  540 value 79.440152
iter  550 value 79.435866
iter  560 value 79.433166
iter  570 value 79.430422
iter  580 value 79.427165
iter  590 value 79.424761
iter  600 value 79.422794
iter  610 value 79.419931
iter  620 value 79.417047
iter  630 value 79.414543
iter  640 value 79.412423
iter  650 value 79.410437
iter  660 value 79.408699
iter  670 value 79.407403
iter  680 value 79.406288
iter  690 value 79.405368
iter  700 value 79.404501
iter  710 value 79.403777
iter  720 value 79.403101
iter  730 value 79.402529
iter  740 value 79.402061
iter  750 value 79.401518
iter  760 value 79.400974
iter  770 value 79.400541
iter  780 value 79.400101
iter  790 value 79.399763
iter  800 value 79.399301
iter  810 value 79.398794
iter  820 value 79.398166
iter  830 value 79.397238
iter  840 value 79.395910
iter  850 value 79.394372
iter  860 value 79.392806
iter  870 value 79.390974
iter  880 value 79.389248
iter  890 value 79.387714
iter  900 value 79.386428
iter  910 value 79.385171
iter  920 value 79.383997
iter  930 value 79.383191
iter  940 value 79.382280
iter  950 value 79.381072
iter  960 value 79.379908
iter  970 value 79.378570
iter  980 value 79.377165
iter  990 value 79.375266
iter 1000 value 79.372790
final  value 79.372536 
stopped after 1001 iterations
Initial stress        : 0.01208
stress after   0 iters: 0.01208
iter   10 value 90.111058
iter   20 value 68.920831
iter   30 value 64.350986
iter   40 value 62.613424
iter   50 value 61.902349
iter   60 value 61.509005
iter   70 value 61.308546
iter   80 value 61.166473
iter   90 value 61.073674
iter  100 value 61.005704
iter  110 value 60.959572
iter  120 value 60.926890
iter  130 value 60.906417
iter  140 value 60.890532
iter  150 value 60.879288
iter  160 value 60.869269
iter  170 value 60.860879
iter  180 value 60.854595
iter  190 value 60.849329
iter  200 value 60.844481
iter  210 value 60.840828
iter  220 value 60.837296
iter  230 value 60.834277
iter  240 value 60.831180
iter  250 value 60.828841
iter  260 value 60.826672
iter  270 value 60.824913
iter  280 value 60.823368
iter  290 value 60.822080
iter  300 value 60.820443
iter  310 value 60.819197
iter  320 value 60.818141
iter  330 value 60.817149
iter  340 value 60.816417
iter  350 value 60.815549
iter  360 value 60.814664
iter  370 value 60.813896
iter  380 value 60.813273
iter  390 value 60.812732
iter  400 value 60.812273
iter  410 value 60.811849
iter  420 value 60.811452
iter  430 value 60.811015
iter  440 value 60.810665
iter  450 value 60.810309
iter  460 value 60.809993
iter  470 value 60.809675
iter  480 value 60.809393
iter  490 value 60.809114
iter  500 value 60.808849
iter  510 value 60.808578
iter  520 value 60.808367
iter  530 value 60.808120
iter  540 value 60.807907
iter  550 value 60.807693
iter  560 value 60.807518
iter  570 value 60.807332
iter  580 value 60.807139
iter  590 value 60.806922
iter  600 value 60.806708
iter  610 value 60.806493
iter  620 value 60.806276
iter  630 value 60.806097
iter  640 value 60.805925
iter  650 value 60.805722
iter  660 value 60.805564
iter  670 value 60.805433
iter  680 value 60.805322
iter  690 value 60.805193
iter  700 value 60.805073
iter  710 value 60.804913
iter  720 value 60.804753
iter  730 value 60.804548
iter  740 value 60.804359
iter  750 value 60.804142
iter  760 value 60.803908
iter  770 value 60.803681
iter  780 value 60.803442
iter  790 value 60.803186
iter  800 value 60.802972
iter  810 value 60.802768
iter  820 value 60.802542
iter  830 value 60.802286
iter  840 value 60.801996
iter  850 value 60.801807
iter  860 value 60.801584
iter  870 value 60.801395
iter  880 value 60.801205
iter  890 value 60.801012
iter  900 value 60.800835
iter  910 value 60.800669
iter  920 value 60.800510
iter  930 value 60.800366
iter  940 value 60.800258
iter  950 value 60.800145
iter  960 value 60.800048
iter  970 value 60.799980
iter  980 value 60.799913
iter  990 value 60.799831
iter 1000 value 60.799746
final  value 60.799737 
stopped after 1001 iterations
iter   10 value 93.987847
iter   20 value 32.507659
iter   30 value 18.676327
iter   40 value 13.901117
iter   50 value 11.929466
iter   60 value 11.030703
iter   70 value 10.569319
iter   80 value 10.283426
iter   90 value 10.078015
iter  100 value 9.943636
iter  110 value 9.848995
iter  120 value 9.771719
iter  130 value 9.715000
iter  140 value 9.670260
iter  150 value 9.635268
iter  160 value 9.607777
iter  170 value 9.583177
iter  180 value 9.561334
iter  190 value 9.541574
iter  200 value 9.522271
iter  210 value 9.508248
iter  220 value 9.495698
iter  230 value 9.484180
iter  240 value 9.473251
iter  250 value 9.462369
iter  260 value 9.452864
iter  270 value 9.444601
iter  280 value 9.437065
iter  290 value 9.429647
iter  300 value 9.422556
iter  310 value 9.416546
iter  320 value 9.410882
iter  330 value 9.406214
iter  340 value 9.400954
iter  350 value 9.396414
iter  360 value 9.392278
iter  370 value 9.388533
iter  380 value 9.384640
iter  390 value 9.381461
iter  400 value 9.378520
iter  410 value 9.375225
iter  420 value 9.372376
iter  430 value 9.369996
iter  440 value 9.367761
iter  450 value 9.365646
iter  460 value 9.363612
iter  470 value 9.361814
iter  480 value 9.360025
iter  490 value 9.358209
iter  500 value 9.356591
iter  510 value 9.354851
iter  520 value 9.353486
iter  530 value 9.352149
iter  540 value 9.350776
iter  550 value 9.349585
iter  560 value 9.348446
iter  570 value 9.347302
iter  580 value 9.346161
iter  590 value 9.344988
iter  600 value 9.343912
iter  610 value 9.342978
iter  620 value 9.342080
iter  630 value 9.341179
iter  640 value 9.340309
iter  650 value 9.339376
iter  660 value 9.338563
iter  670 value 9.337815
iter  680 value 9.337128
iter  690 value 9.336469
iter  700 value 9.335850
iter  710 value 9.335263
iter  720 value 9.334616
iter  730 value 9.333991
iter  740 value 9.333397
iter  750 value 9.332878
iter  760 value 9.332340
iter  770 value 9.331774
iter  780 value 9.331259
iter  790 value 9.330738
iter  800 value 9.330258
iter  810 value 9.329722
iter  820 value 9.329064
iter  830 value 9.328395
iter  840 value 9.327772
iter  850 value 9.326982
iter  860 value 9.326141
iter  870 value 9.325390
iter  880 value 9.324641
iter  890 value 9.323817
iter  900 value 9.322879
iter  910 value 9.322015
iter  920 value 9.320941
iter  930 value 9.319808
iter  940 value 9.318707
iter  950 value 9.317718
iter  960 value 9.316625
iter  970 value 9.315514
iter  980 value 9.314378
iter  990 value 9.313035
iter 1000 value 9.311665
final  value 9.311500 
stopped after 1001 iterations
Initial stress        : 0.00750
stress after   1 iters: 0.00737
iter   10 value 76.056557
iter   20 value 63.434836
iter   30 value 61.570276
iter   40 value 60.878573
iter   50 value 60.541909
iter   60 value 60.358962
iter   70 value 60.269858
iter   80 value 60.216601
iter   90 value 60.182457
iter  100 value 60.159583
iter  110 value 60.144671
iter  120 value 60.131427
iter  130 value 60.119714
iter  140 value 60.112372
iter  150 value 60.105942
iter  160 value 60.100222
iter  170 value 60.095554
iter  180 value 60.092143
iter  190 value 60.089183
iter  200 value 60.086368
iter  210 value 60.084362
iter  220 value 60.082695
iter  230 value 60.081294
iter  240 value 60.080026
iter  250 value 60.079049
iter  260 value 60.078166
iter  270 value 60.077440
iter  280 value 60.076813
iter  290 value 60.076217
iter  300 value 60.075605
iter  310 value 60.075032
iter  320 value 60.074574
iter  330 value 60.074086
iter  340 value 60.073628
iter  350 value 60.073214
iter  360 value 60.072868
iter  370 value 60.072569
iter  380 value 60.072247
iter  390 value 60.071944
iter  400 value 60.071696
iter  410 value 60.071440
iter  420 value 60.071196
iter  430 value 60.070976
iter  440 value 60.070766
iter  450 value 60.070553
iter  460 value 60.070360
iter  470 value 60.070174
iter  480 value 60.069992
iter  490 value 60.069826
iter  500 value 60.069645
iter  510 value 60.069496
iter  520 value 60.069354
iter  530 value 60.069219
iter  540 value 60.069125
iter  550 value 60.069004
iter  560 value 60.068893
iter  570 value 60.068793
iter  580 value 60.068697
iter  590 value 60.068602
iter  600 value 60.068510
iter  610 value 60.068423
iter  620 value 60.068343
iter  630 value 60.068259
iter  640 value 60.068187
iter  650 value 60.068109
iter  660 value 60.068055
iter  670 value 60.067989
iter  680 value 60.067928
iter  690 value 60.067864
iter  700 value 60.067811
iter  710 value 60.067762
iter  720 value 60.067707
iter  730 value 60.067669
iter  740 value 60.067632
iter  750 value 60.067589
iter  760 value 60.067547
iter  770 value 60.067501
iter  780 value 60.067461
iter  790 value 60.067428
iter  800 value 60.067396
iter  810 value 60.067356
iter  820 value 60.067328
iter  830 value 60.067299
iter  840 value 60.067272
iter  850 value 60.067245
iter  860 value 60.067219
iter  870 value 60.067188
iter  880 value 60.067167
iter  890 value 60.067148
iter  900 value 60.067125
iter  910 value 60.067103
iter  920 value 60.067084
iter  930 value 60.067064
iter  940 value 60.067044
iter  950 value 60.067028
iter  960 value 60.067007
iter  970 value 60.066989
iter  980 value 60.066973
iter  990 value 60.066959
iter 1000 value 60.066942
final  value 60.066941 
stopped after 1001 iterations
iter   10 value 59.171571
iter   20 value 17.510960
iter   30 value 8.508536
iter   40 value 5.570668
iter   50 value 4.180655
iter   60 value 3.511969
iter   70 value 3.205928
iter   80 value 3.016826
iter   90 value 2.899066
iter  100 value 2.825574
iter  110 value 2.772954
iter  120 value 2.733537
iter  130 value 2.704694
iter  140 value 2.680982
iter  150 value 2.663163
iter  160 value 2.646997
iter  170 value 2.634993
iter  180 value 2.625126
iter  190 value 2.616654
iter  200 value 2.609508
iter  210 value 2.602977
iter  220 value 2.597315
iter  230 value 2.592247
iter  240 value 2.587458
iter  250 value 2.583513
iter  260 value 2.579839
iter  270 value 2.576642
iter  280 value 2.573482
iter  290 value 2.571027
iter  300 value 2.568626
iter  310 value 2.566357
iter  320 value 2.564102
iter  330 value 2.562115
iter  340 value 2.560065
iter  350 value 2.558339
iter  360 value 2.556707
iter  370 value 2.555225
iter  380 value 2.553611
iter  390 value 2.552184
iter  400 value 2.550660
iter  410 value 2.549406
iter  420 value 2.548128
iter  430 value 2.547057
iter  440 value 2.545940
iter  450 value 2.544920
iter  460 value 2.543885
iter  470 value 2.542927
iter  480 value 2.541994
iter  490 value 2.541137
iter  500 value 2.540385
iter  510 value 2.539580
iter  520 value 2.538779
iter  530 value 2.537959
iter  540 value 2.537278
iter  550 value 2.536635
iter  560 value 2.535965
iter  570 value 2.535383
iter  580 value 2.534861
iter  590 value 2.534284
iter  600 value 2.533737
iter  610 value 2.533229
iter  620 value 2.532800
iter  630 value 2.532437
iter  640 value 2.532031
iter  650 value 2.531656
iter  660 value 2.531318
iter  670 value 2.530948
iter  680 value 2.530636
iter  690 value 2.530320
iter  700 value 2.530055
iter  710 value 2.529806
iter  720 value 2.529503
iter  730 value 2.529258
iter  740 value 2.529021
iter  750 value 2.528809
iter  760 value 2.528551
iter  770 value 2.528310
iter  780 value 2.527987
iter  790 value 2.527671
iter  800 value 2.527360
iter  810 value 2.527114
iter  820 value 2.526888
iter  830 value 2.526677
iter  840 value 2.526482
iter  850 value 2.526303
iter  860 value 2.526113
iter  870 value 2.525902
iter  880 value 2.525680
iter  890 value 2.525438
iter  900 value 2.525239
iter  910 value 2.525041
iter  920 value 2.524836
iter  930 value 2.524628
iter  940 value 2.524403
iter  950 value 2.524213
iter  960 value 2.524021
iter  970 value 2.523816
iter  980 value 2.523628
iter  990 value 2.523464
iter 1000 value 2.523309
final  value 2.523286 
stopped after 1001 iterations
Initial stress        : 0.00747
stress after   0 iters: 0.00747
```

In [29]:

```
#MSE
result <- rbind(MSE_MDS, MSE_H1, MSE_H2)
colnames(result) <- Ms
rownames(result) <- c("MDS","H1", "H2")
result
```

A matrix: 3 × 4 of type dbl

|  | 5 | 10 | 20 | 30 |
| --- | --- | --- | --- | --- |
| MDS | 0.22843070 | 0.045991711 | 0.0248239345 | 0.0252232623 |
| H1 | 0.01579753 | 0.004016482 | 0.0030248625 | 0.0029884050 |
| H2 | 0.02509713 | 0.003704609 | 0.0004400514 | 0.0001204638 |

In [30]:

```
#Angle
yrange <- c(0, max(max(angle_H1/(2*pi)*360),max(angle_H2/(2*pi)*360)))
par(cex=1.3)
tmp <- angle_MDS/(2*pi)*360
tmp2 <- list("M=5"=tmp[,1],"M=10"=tmp[,2],"M=20"=tmp[,3],"M=30"=tmp[,4])
boxplot(tmp2, ylim=yrange, ylab="Angles in degrees", main="MDS", cex.lab=1.4, cex.axis=1.4, cex.main=3)

tmp <- angle_H1/(2*pi)*360
tmp2 <- list("M=5"=tmp[,1],"M=10"=tmp[,2],"M=20"=tmp[,3],"M=30"=tmp[,4])
boxplot(tmp2, ylim=yrange, ylab="Angles in degrees", main="H1", cex.lab=1.4, cex.axis=1.4, cex.main=3)

tmp <- angle_H2/(2*pi)*360
tmp2 <- list("M=5"=tmp[,1],"M=10"=tmp[,2],"M=20"=tmp[,3],"M=30"=tmp[,4])
boxplot(tmp2, ylim=yrange, ylab="Angles in degrees", main="H2", cex.lab=1.4, cex.axis=1.4, cex.main=3)
```

In [ ]:
